# Neuroblastoma-associated ALK variants have distinct cellular and biochemical activities

**DOI:** 10.1101/2025.09.25.677354

**Authors:** Anna M. Wulf, Stuart Lutimba, Dylan Cameron, Jade Desjardins, Tanya J. Shaw, Leone Rossetti, Mohammed A. Mansour, Karen J. Liu

## Abstract

Mutations in anaplastic lymphoma kinase (ALK) are associated with high-risk neuroblastoma, a childhood cancer arising in trunk neural crest cells. The role of ALK in undifferentiated NC is still unknown; however, the presence of activating mutations in ALK correlates with migratory and invasive cell behaviours in neuroblastoma cell lines. Here, we show the functional consequences of ALK overexpression on neural crest cells, by comparing wildtype ALK (ALK^WT^) protein to ALK gain-of-function variants ALK^F1174L^ and ALK^R1275Q^. Elevated ALK activity, independent of mutational status leads to increased migration velocity and loss of directionality, while ALK^F1174L^ overexpression presents additional effects on cytoskeletal protrusions. These results correlate with increased binding of ALK^F1174L^ to GSK3, which has previously been shown to regulate cytoskeletal dynamics in neural crest cells. Further, molecular dynamics simulations of the ALK-GSK3 complex show high flexibility, suggestive of enhanced allosteric regulation. Together, our data show that activating mutations in ALK drive migratory changes in trunk NC cells, potentially mediated by its novel interacting partner GSK3.

**Significance Statement:** Anaplastic lymphoma kinase (ALK)-associated neuroblastoma phenotypes arise in the neural crest lineage, but the effects on neural crest cell behaviours are not well-studied. Here, we use primary mouse neural crest cells to study the functional relevance of ALK-activating mutations on cell migration, which we then biochemically link to interactions with GSK3-isoforms. We also directly compare the activities of wild-type ALK with two gain-of-function variants associated with metastatic disease, ALK^F1174L^ and ALKR^1275Q^, and find that ALK^F1174L^ causes increased cytoskeletal protrusions and GSK3 binding. Our study provides insights into a signalling pathway in neural crest that drives cytoskeletal rearrangements and pathological migration in neuroblastoma.

## Introduction

Neuroblastoma is a childhood cancer derived from trunk neural crest cells, with approximately 40% of cases diagnosed before the age of one^1^. In the UK, roughly 95 children a year are newly diagnosed with neuroblastoma, accounting for a total of 6% of all childhood cancers^2^. However, 5-year survival rates in neuroblastoma cases are the lowest among all children with cancer in the UK^2^, in part due to almost half of cases diagnosed at late stages^3^.

Neuroblastoma is a highly heterogeneous tumour composed of undifferentiated progenitor cells^4,5^. Although numerous genetic alterations have been identified, it remains unclear at which developmental stage these cells become malignant^7^. Mutations in genes important during neural crest development lead to tumours hijacking embryonic signalling pathways to sustain proliferation, block differentiation, and ultimately metastasize throughout the body^8–10^. Primary tumours can arise anywhere along the sympathetic nervous system, with approximately half originating in the adrenal medulla^6^.

The tyrosine kinase receptor *anaplastic lymphoma kinase* (*ALK*) is the most frequently mutated gene in neuroblastoma. Mutations within *ALK* are found in an estimated 11% of newly diagnosed cases, with enrichment in stage 3 or stage 4^11^. In neuroblastoma, ALK protein is known to direct cell migration and invasion through extracellular matrix remodelling^12,13^, while cleavage of its extracellular domain influences epithelial to mesenchymal transition^14^. In neural crest cells, expression of *ALK* or its orthologue *LTK* is spatiotemporally linked to migration and delamination^15,16^. Overexpression of constitutively active *ALK* variants leads to prolonged sympathetic neuron progenitor proliferation^17,18^ and subsequent blockage of downstream differentiation^19,20^. However, our mechanistic understanding of how ALK regulates these cell behaviours is limited. Most downstream targets have been identified in the context of other diseases, or with oncogenic ALK-fusion proteins, while the ALK variants associated with neuroblastoma are rarely considered (reviewed in^21^).

In neuroblastoma, 85% of *ALK* mutations are found at three distinct positions in its kinase domain: F1174, F1245, R1275^11^. Almost half of the neuroblastoma-associated ALK mutations are located at R1275, whereas F1174 and F1245 are found in smaller proportions at 30% and 12% of all cases, respectively^11^. *ALK* mutations are associated with refractory and metastatic cases^11^; cases with additional MYCN amplifications are classified as high-risk tumours with poor outcomes^22^. Interestingly, even though most of the mutations lead to ligand independent constitutive kinase activity^21^, they are present in distinct subsets of neuroblastoma cases. R1275 mutations can be found in familial and sporadic tumours, whereas F1174 and F1245 variants are mostly found in sporadic cases^23^. In relapses, *ALK* mutations are significantly increased, with the F1174 and R1275 variants being prevalent^11^. However, we currently lack a clear understanding of altered cell behaviour induced by these mutations.

In this study, we directly compare wildtype ALK activity (ALK^WT^) to the two most prevalent neuroblastoma-associated ALK mutations, ALK^F1174L^ and ALK^R1275Q^. Overexpression of either ALK variant, but not ALK^WT^, leads to loss of contact inhibition of proliferation, phenocopying neuroblastoma cell lines. Using primary mouse neural crest cells, we observe increased migration speed, loss of directionality and extended protrusions induced by ALK^F1174L^ overexpression. In contrast, ALK^WT^ or ALK^R1275Q^-overexpressing cells had mild phenotypes limited to a loss of directionality. We previously hypothesised that ALK can influence neural crest cell behaviours by interacting with glycogen synthase kinase 3 (GSK3), a serine/threonine kinase involved in neural crest migration^15,24^. Here, we confirm this interaction and demonstrate that it requires not only a functioning ALK kinase domain but is notably increased with ALK^F1174L^ compared to ALK^WT^ or ALK^R1275Q^. *In silico* modelling suggests that these differences are not due to conformational changes, but rather through changes in protein dynamics introduced by the amino acid substitution. These data are important for understanding ALK-variant induced cell signalling relevant to neuroblastoma pathology.

## Results

### Mutant active ALK leads to loss of contact inhibition of proliferation, a hallmark of cancer

Loss of contact inhibition of proliferation is commonly seen in cancer, and is connected to the increased ability of cancer cells to invade surrounding tissues^25^. When cultured, these cells lose the ability to sense high density environments resulting in sustained proliferation. Architecturally, cells change from a 2D monolayer, developing multiple cell layers or cell clumps^26^. We examined this phenotype in four neuroblastoma cell lines, comparing ALK^WT^ or ALK^F1174L^-positive genotypes, with or without MYCN amplification (Figure 1A). The NBLS cell line (normal ALK^WT^/MYCN^WT^) showed a typical monolayer culture. In contrast, we observed loss of contact inhibition particularly in ALK^F1174L^ carrying SH-SY5Y cell line, with cell clumps presenting in confluent areas of the dish (Figure 1A). Milder phenotypes were observed in the other cell lines IMR-5 (MYCN^AMP^) and in Kelly (ALK^F1174L^/MYCN^AMP^). We confirmed increased ALK activity (phosphorylated ALK) in SH-SY5Y and Kelly cells compared to ALK^WT^ cell lines (NBLS and IMR-5, Figure 1B).

**Figure 1:**
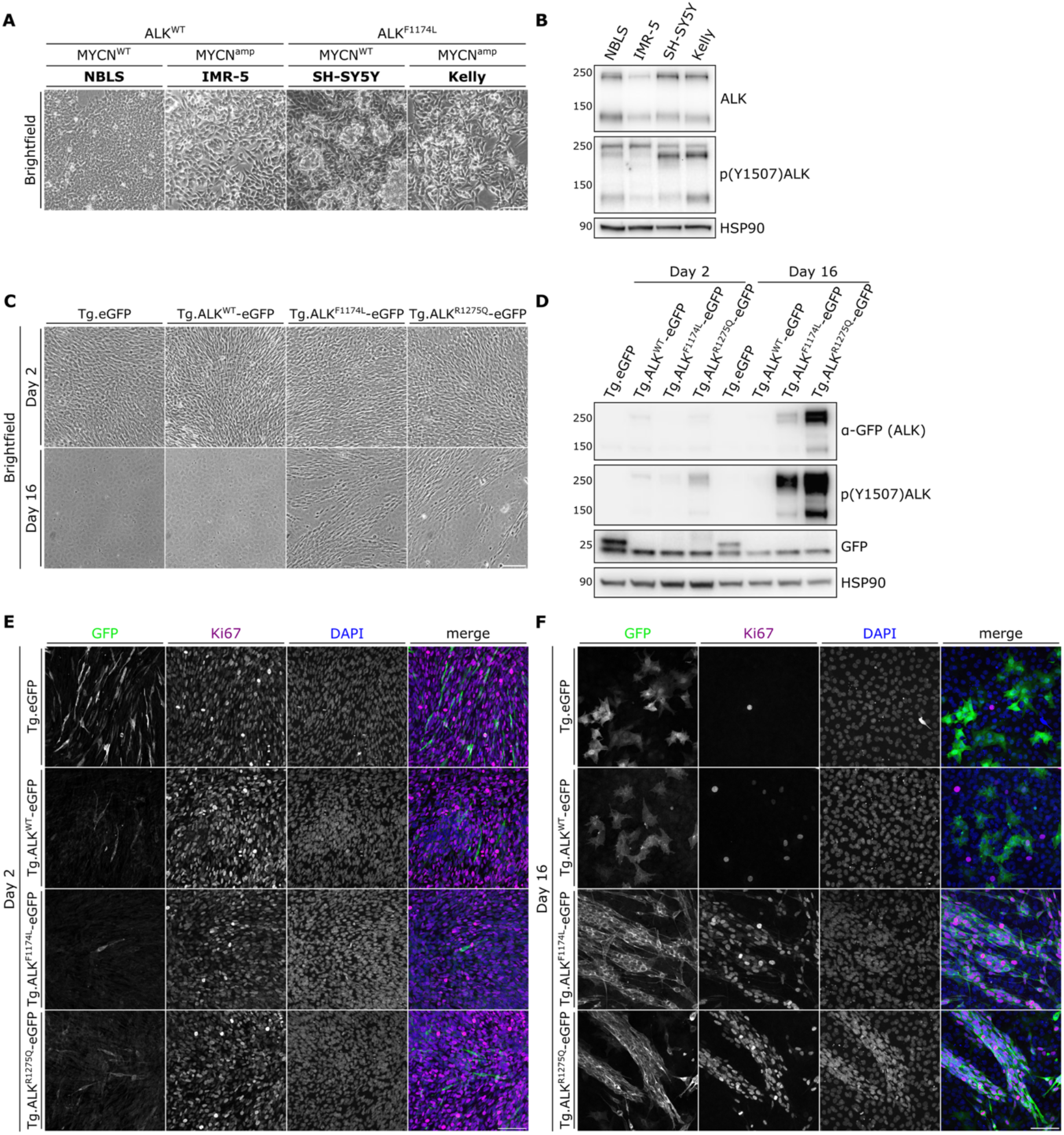
ALK^F1174L^ and ALK^R1275Q^ induce loss of contact inhibition of proliferation. (A) Brightfield images of neuroblastoma cell lines with different ALK and MYCN status at confluency. (B) Western blot of neuroblastoma cell lines probed with anti-ALK and anti-p(Y1507)ALK (active ALK). ALK is present as two bands. HSP90 was used as loading control. (C-F) Transgenic O9-1 cells were cultured for either two days or 16 days, followed by fixation for (C) brightfield imaging and (E-F) immunofluorescence imaging, or lysed for (D) Western blot analysis. Blot was probed with anti-GFP (eGFP and ALK-eGFP) and anti-p(Y1507)ALK (active ALK). HSP90 was used as loading control. (E-F) Immunostaining of transgenic O9-1 cells at day 2 (E) or day 16 (F), stained for GFP (green, eGFP or eGFP-tagged ALK), Ki67 (magenta, proliferation marker), and DAPI (blue, nuclei). (F) Day 16 ALK^F1174L^ and ALK^R1275Q^ retain proliferation and form 3D multilayer structures. Scale bar: 100μm.

While cell lines have widely been used to study neuroblastoma behaviour, they often carry additional mutations, making it difficult to attribute specific phenotypes to single mutations. To overcome this, we generated transgenic murine neural crest cells (O9-1)^27^ overexpressing either eGFP, ALK^WT^-eGFP, ALK^F1174L^-eGFP or ALK^R1275Q^-eGFP. These lines were then challenged by culturing to confluency, where cells stop proliferating to generate a monolayer culture (see Figure 1C, Day 16 Tg.eGFP and Tg.ALK^WT^-eGFP). Conversely, both Tg.ALK^F1174L^-eGFP and Tg.ALK^R1275Q^-eGFP cultures develop 3D multi-layer cell structures, where individual cells retain elongated morphology as seen in the early culturing conditions (Figure 1C). Western blotting analysis shows significant increase in expression of Tg.ALK^F1174L^-eGFP or Tg.ALK^R1275Q^-eGFP at late stages of culture (Figure 1D), suggesting that variant ALK-expressing cells are the main contributors to the overgrowth. In confirmation, we assessed proliferation at both early and late stages of culturing. At early stages most cells were proliferating (Ki67 positive), but very few transgenic ALK-positive cells were detected (Figure 1E). By day 16, when cells were confluent, we found that proliferation is only retained in Tg.ALK^F1174L-^ eGFP and Tg.ALK^R1275Q^-eGFP positive cells (Figure 1F). Additionally, the 3D structures are solely made up of eGFP-positive cells, suggesting the cause of loss of contact inhibition is mutant active ALK.

Together, we show that loss of contact inhibition of proliferation, an *in vitro* phenotype of cancer cells, can be induced by overexpression of either ALK^F1174L^ or ALK^R1275Q^, whereas overexpression of ALK^WT^ is not sufficient.

### Constitutively active ALK variants perturb trunk neural crest cell migration and cell morphology

ALK is often described to promote neuroblastoma migration and invasion^14^ but direct consequences on cell migration are unknown. Additionally, ALK-dependent loss of contact inhibition of proliferation suggests an invasive and migratory cell behaviour. To identify cell pathologies driven by ALK, we turned to mouse embryos, which allow us to isolate primary neural crest cells at the relevant axial level. We dissected the neural plate border adjacent to somites 18-21 from embryonic day 9.5 (E9.5) mouse embryos. After 24 hours in culture, neural crest cells emigrate in a halo surrounding the neural plate border (Figure 1A). We then transfected these cultures with control eGFP, ALK^WT^-eGFP, ALK^F1174L^-eGFP, or ALK^R1275Q^-eGFP and performed live-imaging for 24 hours followed by fixation and immunofluorescence imaging.

First, we evaluated the overall morphological changes resulting from ALK overexpression (Figure 2B-D). eGFP and ALK^WT^ overexpressing cells are circular, spread out and display filopodial and lamellipodial protrusions at their leading edge (Figure 2B-D). Cells overexpressing ALK^F1174L^, and to a lesser extent ALK^R1275Q^, displayed an elongated morphology with decreased cell circularity (Figure 2B/D), whereas cell area remains unchanged (Figure 2C). Notably, cells overexpressing ALK^F1174L^-eGFP display extended protrusions (Figure 2B).

**Figure 2:**
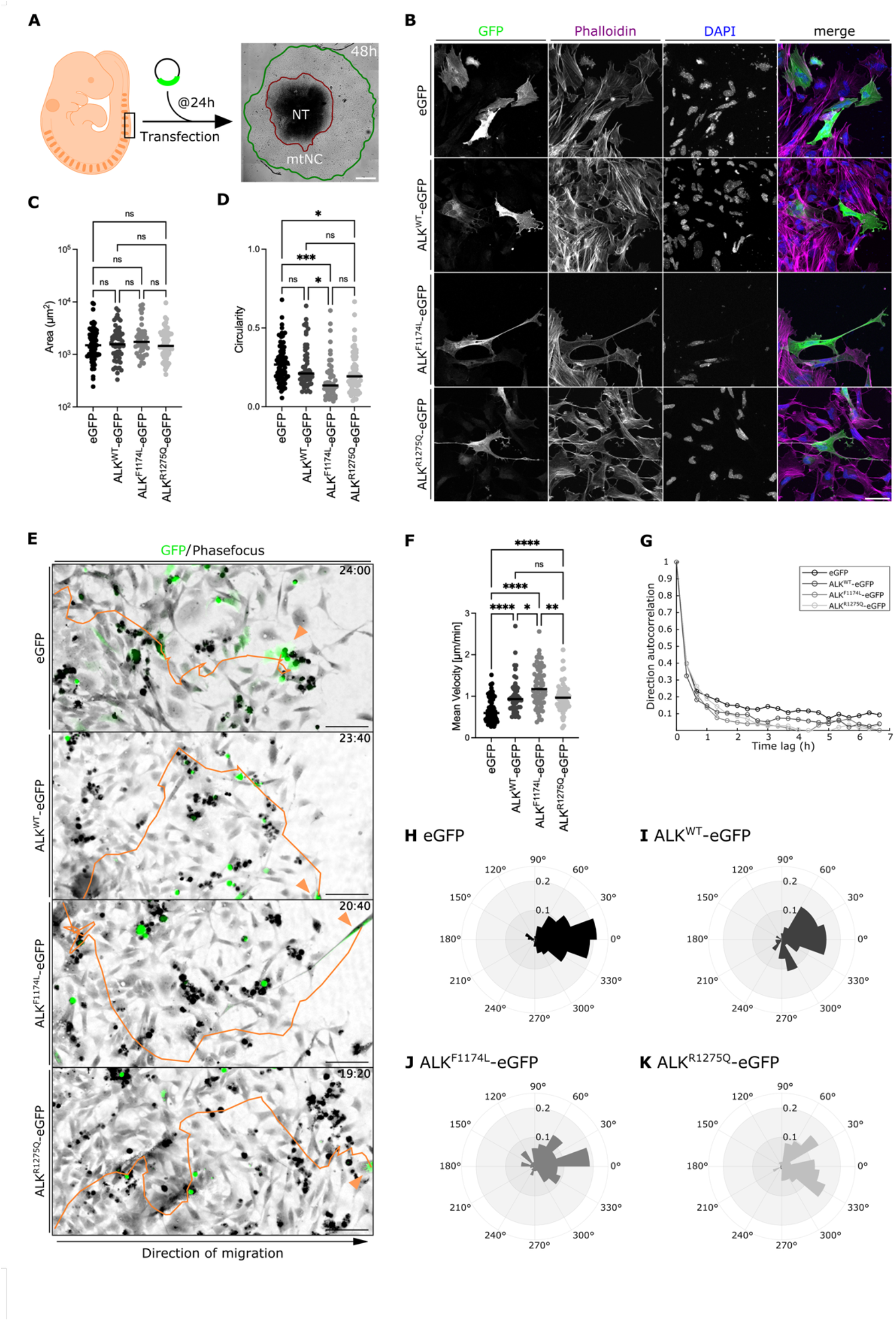
ALK overexpression in primary trunk neural crest cells leads to increased migration and loss of directionality. (A) For primary trunk neural crest cell explants, the neural plate border of E9.5 mouse embryos was dissected (black box) and cultured. Transfections were performed at 24h post-dissection and time lapse imaging for 24 hours post-transfection. Explants were fixed after time lapse imaging for immunostaining. (B) Immunostaining for GFP (green, eGFP and eGFP-tagged ALK), Phalloidin (magenta, actin filaments), and DAPI (blue, nuclei) of trunk neural crest explants 48h post-transfection. Scale bar: 50μm. (C) Quantification of cell area (μm^2^) and cell circularity of GFP positive cells. (E-K) Time lapse imaging for 24h every 20min. For quantification, GFP positive cells were manually tracked using ImageJ. (E) Example tracks per condition (orange lines). (F) Quantification of mean velocity (μm/min) of GFP positive cells. (G-K) x- and y-coordinates were used to generate (G) directional autocorrelation and (H-K) rose plots. (E-K) Shown are representative examples of a set of 3 biological replicates (n=3), each data point represents one cell. *p<0.05, **p<0.001, ***p<0.0001, ****p<0.00001 One-Way ANOVA. See Supplementary Movie S1.

We assessed the migratory behaviour of these cells by manually tracking GFP positive cells (Figure 2E, Supplementary Movie S1). Overall, overexpression of ALK leads to increased migration when compared to the eGFP controls with the most increase in ALK^F1174L^ (Figure 2F). Migration directionality was visualised using rose plots and direction autocorrelation^28^. eGFP-overexpressing cells exhibited a unimodal distribution centred at 0°, indicative of persistent, directed migration (Figure 2F). In contrast, ALK-overexpressing cells displayed broader angular dispersion and lacked a dominant orientation (Figure 2I-K). Direction autocorrelation was consistently higher in eGFP cells, further supporting their enhanced migratory persistence (Figure 2G).

In summary, ALK overexpression in primary trunk neural crest cells leads to increased velocity and loss of directionality. However, only the overexpression of ALK^F1174L^ induces significant changes in cell morphology leading to elongated cells and extended protrusions.

### GSK3 interacts with ALK in a variant-specific manner

We have recently shown that the serine-threonine kinase GSK3 plays an important role during lamellipodium formation in neural crest by controlling actin dynamics and the stability of focal adhesions^24^. This is of particular interest, as *Alk* and *Gsk3* expression is spatiotemporally linked in migrating neural crest cells^15^. Additionally, GSK3 contains multiple ALK-phosphorylation motifs, which raises the possibility that ALK may act via GSK3. To assess a physical interaction, we co-expressed GSK3α/β-Flag and ALK^WT^ in HEK293T cells (Figure 3A). We found that either GSK3α or GSK3β can be immunoprecipitated with ALK^WT^ (Figure 3A’).

**Figure 3:**
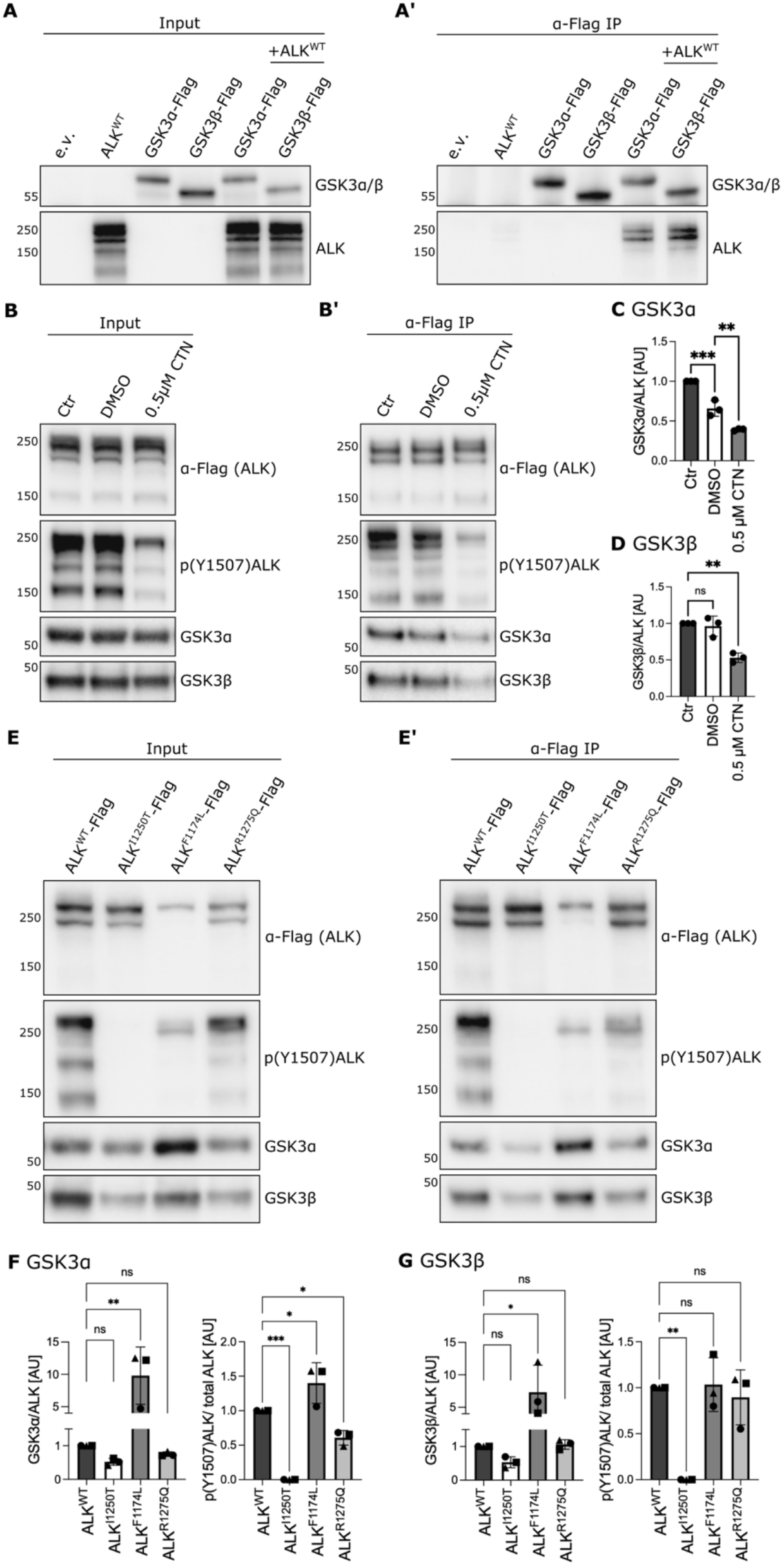
Differential interactions between ALK variants with GSK3 isoforms. (A-A’) Immunoprecipitation of GSK3α/β-Flag with ALK^WT^, probed with anti-Flag (GSK3α/β) and anti-ALK. n=3 (B-B’) Immunoprecipitation of ALK-Flag with either GSK3α or GSK3β, 24h after cells were treated with 0.5μM crizotinib (CTN). DMSO was used as controls. Probed with anti-Flag (ALK), anti-p(Y1507)ALK (active ALK), anti-GSK3α, and anti-GSK3β. (C-D) Quantification of B-B’ using ImageJ. (E-G) Immunoprecipitation of ALK^WT^-Flag, ALK^I1250T^-Flag (kinase-dead), ALK^F1174L^-Flag, or ALK^R1275Q^-Flag with either GSK3α or GSK3β, probed with anti-Flag (ALK), anti-p(Y1507)ALK (active ALK), anti-GSK3α, and anti-GSK3β. (F-G) Quantification of E-E’ using ImageJ. *p<0.05, **p<0.001, ***p<0.0001, One-Way ANOVA. Each data point represents one biological repeat (n=3), error bars are standard deviation,

Next, we tested whether this interaction is dependent on the kinase activity of ALK. Here, we co-expressed ALK^WT^-Flag with either GSK3α or GSK3β and treated cells for 24h with the ALK inhibitor crizotinib (CTN) (Figure 3B). We found that interactions between ALK^WT^-Flag and GSK3α or GSK3β were significantly reduced when ALK activity was inhibited (Figure 3B-D). To exclude potential steric interference induced by crizotinib, we also performed immunoprecipitations with the kinase-dead mutant ALK^I1250T^-Flag (Figure 3E-G). We show a consistent decrease in binding between kinase-dead ALK^I1250T-^Flag and GSK3α or GSK3β (Figure 3E-G). Together, this suggests that a functional ALK kinase domain is a requirement for this interaction.

Finally, we tested the interaction between GSK3α/β with neuroblastoma associated mutations ALK^F1174L^ and ALK^R1275Q^ (Figure 3E-G). To our surprise, GSK3α and GSK3β show increased affinity to ALK^F1174L^-Flag, whereas ALK^R1275Q^-Flag co-precipitated GSK3α or GSK3β at similar levels as seen with ALK^WT^-Flag (Figure 3F-G). Interestingly, high ALK^F1174L^ activity, assessed by phospho-tyrosine ALK, correlated with increased interaction with GSK3α (Figure 3F). This correlation was not seen with GSK3β (Figure 3G).

Together, we demonstrate that ALK and GSK3α/β interact and that this interaction requires a functional ALK kinase domain. As kinase activity cannot fully account for these differences, we hypothesise that conformational changes and protein dynamics may may underlie isoform-specific differences in binding interactions and activity.

### Structural modeling of ALK mutants in complex with GSK3 isoforms

To gain mechanistic insight into ALK-GSK3α/β interactions, we conducted molecular dynamics simulations to evaluate how ALK variants and GSK3 isoforms affect the stability, conformational flexibility, and interaction configurations of ALK-GSK3 complexes in near-physiological environments.

We repurposed experimental crystal structures of the ALK kinase domain (Q9UM73) and both GSK3 isoforms (GSK3α (P49840) and GSK3β (P49841) from the Uniprot database. Since several regions in GSK3α/β were unresolved in Uniprot, we utilized a modelling approach to model and refine their structures based on sequence data and evolutionary information. Using AlphaFold, we reconstructed full-length models of the ALK–GSK3α/β complexes, focusing primarily on the kinase domains, as they are essential for catalytic activity and protein–protein interactions that directly influence the functionality of the complex. The reconstructed models of models the kinase domains for ALK, GSK3α and GSK3β were obtained with a high-confidence score (pLDDT > 90) for the kinase core and moderate confidence for more flexible terminal regions. These models were further stereo-chemically validated using Ramachandran plot analysis, with particular attention to the N- and C-terminal segments. Using the High Ambiguity Driven Biomolecular Docking (HADDOCK) platform, we were able to model the interaction between ALK^WT^ (residue 1105-1377) and GSK3α (residues 82 to 457) (Figure 4A) or GSK3β (residue 1-403) (Figure 4B) with protein-protein interaction distances of ≤3Å, highlighting close contact.

**Figure 4:**
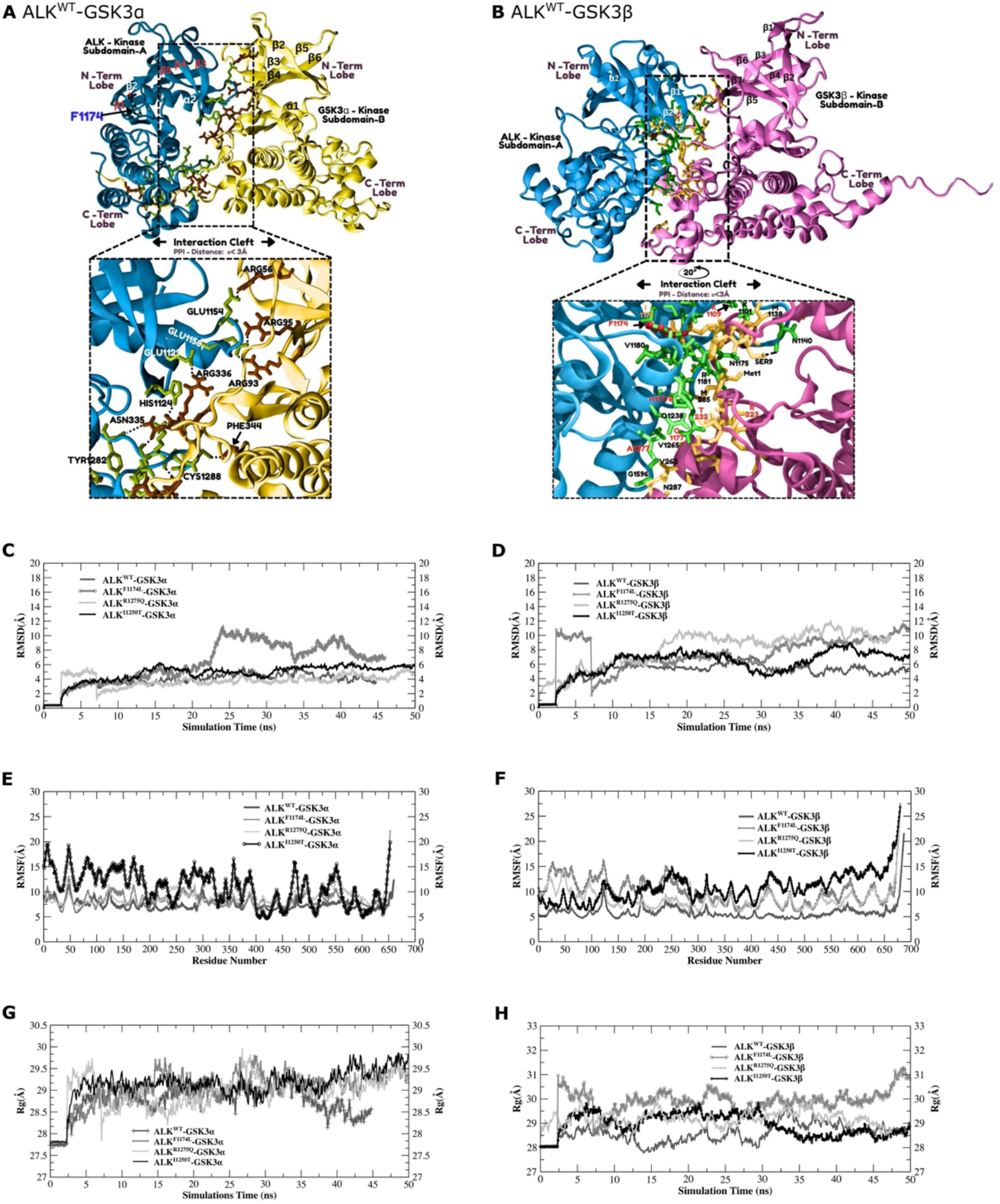
Structural comparison of the protein–protein interfaces between ALK and GSK3α/β derived from 50 ns molecular dynamics simulations. (A-B) Complex formation between kinase domain of ALK^WT^ with GSK3α (A, yellow) or GSK3β (B, pink). Enlarged interaction cleft (distance <3 Å) reveals side-chain contacts that stabilize the complex: ALK residues are shown as green sticks, and GSK3 residues in brown (α) and yellow (β). Panel B inset is rotated 20° from above view to illustrate binding geometry. Root Mean Square Deviation (RMSD) showing complex dynamics (C-D), Root Mean Square Fluctuation (RMSF) showing flexibility (E/F), and Radius of Gyration (Rg) analysis from 50 ns molecular dynamics simulations of GSK3α (C/E/G) or GSK3β (D/F/H) complexing with either ALK^WT^, ALK^F1174L^ (mutant active), ALK^R1275Q^ (mutant active), or ALK^I1250T^ (kinase-dead). See detailed Supplementary Figure S1 and Supplementary Movie S2-S3 for dynamic simulations.

The ALK-GSK3α interaction is primarily facilitated through polar or charged amino acids (Glu, Arg, His) with contributions through hydrogen bonds, hydrophobic interaction, and possibly π-π stacking via F334 and T1282. On the other hand, the interaction between ALK^WT^ and GSK3β is facilitated through flexibility of the interface and hydrophobic contacts (F1174, V1180) with interacting amino acids for ALK and for GSK3β noted (Figure 4B). Although ALK interacts with GSK3α and GSK3β in slightly different ways, their binding pockets are largely similar. The two isoforms have very similar sequences, but small shifts in their kinase lobes, especially the N-terminus, may influence their specificity and interaction strength with partners like ALK.

### More dynamic, less stable complex formation between ALK variants and GSK3

We then used UCSF Chimera to introduce specific ALK mutations: F1174L, R1275Q, and I1250T. The interaction between ALK^F1174L^-GSK3α is mainly facilitated by hydrophobic and polar interaction (Supplementary Figure S1A), where arginine-rich patches are known to stabilize transient electrostatic contacts. For the ALK^F1174L^-GSK3β complex, relevant small polar and basic residues (T8, R6, N287, K292) at the interface highlight the involvement of flexible regions within GSK3β for the interaction (Supplementary Figure S1D). We observed a more dynamic and less stable complex formation with ALK variants (Figure 4C/D), calculated by Root Mean Square Deviation (RMSD) of the protein backbone over a 50 ns simulation trajectory. ALK^F1174L^ exhibited the highest average RMSD when interacting with either isoform (GSK3α: 6.26 ± 2.76 Å; GSK3β: 7.39 ± 2.46 Å) (Figure 4C/D). This significant increase in structural deviation, indicates a more dynamic and less stable complex conformation compared to ALK^WT^ (GSK3α: 3.82 ± 1.07 Å; GSK3β: 5.06 ± 1.30 Å).

We next probed local flexibility and structural compactness measured using Root Mean Square Fluctuation (RMSF) (Figure 4E/F) and Radius of Gyration (Rg) analyses respectively. As with RMSD, ALK^F1174L^ variant showed the most significant differences, with substantial fluctuations across the complex, particularly in concert with the GSK3β isoform (9.99 ± 2.62 Å vs. WT 5.88 ± 1.50 Å) (Figure 4F). This pronounced flexibility translated into a measurable expansion of the complex (Figure 4G/H, Supplementary Movie S2/S3). The Rg analysis confirmed that ALK^F1174L^ yielded the most expanded conformation for the GSK3β complex (29.86 ± 0.59 Å vs. WT 28.58 ± 0.37 Å) (Figure 4H). A similar, though less dramatic, expansion trend was observed for ALK^F1174L^ in the GSK3α complex (29.04 ± 0.41 Å vs. WT 28.78 ± 0.36 Å) (Figure 4G).

Interestingly, the interaction of ALK^R1275Q^-GSK3α/β partly retains a more ALK^WT^-like interaction, particularly in complex with GSK3β. The ALK^R1275Q^-GSK3α complex retains several stabilizing salt-bridges (GSK3α: E59, R336, K114; ALK^R1275Q^: E1340, T1343) though showing subtle alteration of charge and hydrogen-bonds (Supplementary Figure S1B). Interestingly, RMSD shows a remarkably stable ALK^R1275Q^-GSK3α complex (3.70 ± 1.02 Å), mirroring that of ALK^WT^-GSK3α (Figure 4C). In contrast, ALK^R1275Q^-GSK3β complex is calculated to be destabilized (8.19 ± 2.47 Å) (Figure 4D), with interaction facilitated by a mixture of aromatic and charged residues leading to flexible yet functionally capable interface (Supplementary Figure S1E). The preservation of several positively charged residues and exposure of mutant side chains is consistent with the slightly enhanced but less robust interaction models for ALK^R1275Q^.

When analysing the residue flexibility (RMSF), it is notable that kinase-dead ALK^I1250T^ complexes experience the highest per-residue flexibility (Figure 4E/F). For ALK^I1250T^-GSK3α, increased involvement of flexible termini appears to disrupt hydrogen bonds and hydrophobic networks, likely leading to instability and loss of proper confirmational coupling ^29^. When in complex with GSK3β, a suite of contacts illustrates expanded dynamics and instability (Figure 4E/F). The dispersal of interactions across the GSK3β sequence may reflect compensatory stabilization due to the loss of canonical interface geometry.

Through molecular modelling approaches, we reveal specific residue-level changes at the ALK-GSK3 interface linked to ALK mutations (extended results in supplementary supporting text). We find that ALK kinase activity is required, as I1250T kinase-dead mutation disrupts functional complex geometry. Most exciting, our analysis demonstrates that the ALK^F1174L^ facilitates a plasticity-enhancing interface with increased complex flexibility with both GSK3α and GSK3β, whereas ALK^R1275Q^ only has that with GSK3β, possibly elucidating a key mechanistic basis for differential signalling in neuroblastoma disease development.

## Discussion

Here, we report on the cellular and biochemical functions of ALK, with a focus on the two most common neuroblastoma mutations, ALK^F1174L^ and ALK^R1275Q^. By comparing wild-type and missense variants of *ALK* using primary neural crest cells and biochemical assays, we have identified distinct cell behaviours and biochemical associations. Our data suggest that simply increasing ALK-signalling levels by overexpressing ALK^WT^, is not sufficient to elicit cellular pathology, and furthermore, that not all ALK gain-of-function variants are equivalent. Notably, we find that ALK^F1174L^ induces the most extreme cellular changes and has the highest binding affinity to GSK3 proteins, a known regulator of the cytoskeleton^24,30^. Altogether, our findings raise the possibility that ALK-variants may act to shift the balance of GSK3 activity in neural crest cells, underlying pathological cell behaviours.

The comparison between mutant *ALK* variants is limited in literature. In neuroblastoma, *ALK*^*F1174L*^ is most frequently found at diagnosis, whereas *ALK*^*R1275Q*^ is predominantly found in relapses^11^. Our most remarkable results were the comparison between the two neuroblastoma associated mutations, ALK^F1174L^ and ALK^R1275Q^. Using primary trunk neural crest explants, we report cellular differences induced by ALK^F1174L^ or ALK^R1275Q^. Overall, the ALK^F1174L^ cells migrated quicker and with a significantly elongated cell body, while ALK^R1275Q^ cells were equivalent to ALK^WT^ expressing cells. This correlates with increased affinity of ALK^F1174L^ to GSK3. In comparison, binding of ALK^WT^ or ALK^R1275Q^ to GSK3 was comparable. Interestingly, molecular dynamics simulations revealed global destabilization of ALK^F1174L^-GSK3, characterized by high backbone deviation, increased local flexibility, and a more expanded molecular radius. We propose that this enhanced flexibility could indicate sampling more favourable conformations for high-affinity binding, potentially allowing deeper protein-protein engagement or alternative interaction modes. This would be consistent with recent literature, demonstrating that increased protein flexibility reflected by higher RMSD can foster alternative, functionally relevant, conformational states, ultimately enhancing allosteric regulation and enzymatic output rather than merely weakening binding affinity^32^. The dramatic isoform-specific destabilization of the ALK^R1275Q^-GSK3β complex is in stark contrast to the stable ALK^WT^-like interaction with GSK3α, suggesting that the functional and structural consequences of this mutation are highly context-dependent.

Interestingly, *ALK*^*F1174L*^ is exclusively found as a sporadic mutation, whereas *ALK*^*R1275Q*^ is found as a germline mutation^23^. In support, Stalin and colleagues used transgenic animals to show that *ALK*^*F1174L*^ overexpression in *Sox10+* cells leads to 100% lethality by E12.5^20^, likely due to ALK^F1174L^ priming drastic developmental defects. When ALK^F1174L^ expression is induced using drivers expressed after neural crest induction, such as *tyrosine hydroxylase (TH)*, the mice generally fail to form tumours^31^. Both models preclude the study of the consequences of ALK mutation in migrating neural crest. Our studies focusing on these neural crest migration stages may explain the severity of early *ALK*^*F1174L*^ overexpression *in vivo*.

In our prior studies, inhibition or loss of GSK3 decreased neural crest cell migration, which was phenocopied when ALK was inhibited^15^. Further, GSK3 regulates the actin cytoskeleton by phosphorylating the actin effector protein lamellipodin^24^. In this study, we show that ALK-GSK3 interactions require ALK activity supporting the hypothesis that ALK acts in concert with GSK3. This could be a direct effect, via phosphorylation, or indirect effect, e.g., via recruitment to the plasma membrane, promoting migration. Further modelling of these interactions will help us define cellular subfunctions of both ALK and GSK3 in neural crest, separating proliferative and migratory functions from cell fate specification (e.g. sympathoadrenal differentiation). We may also be able to distinguish between ALK-variant associated pathologies in neuroblastoma and in other cancers which will aid our capacity for targeting distinct tumour phenotypes for therapeutic specificity.

## Materials and Methods

### Cell lines

The following cell lines were used: HEK293T cells (ThermoFisher #R70007) for protein biochemistry and neuroblastoma cell lines NBLS (DMSZ ACC 656), IMR-5 (gift from Martin Eilers, Würzburg, Germany), SH-SY5Y (DMSZ ACC 209), and Kelly (DMSZ ACC 355). All cell lines were cultured according to conventional methods. The mouse cranial neural crest cell line O9-1 was cultured as described in Nguyen et al., 2018^33^, with the following modification: cells were plated onto 1μg/ml fibronectin pre-coated plastic well plates.

### Primary trunk neural crest explants

Animal work was approved by King’s College London Animal Welfare and Ethical Review Body (AWERB) and performed at King’s College London in accordance with the UK Home Office Project Licenses PP1672528 (T. Shaw) and PP3807977 (T. Shaw). After mating, the observation of a vaginal plug was considered embryonic day 0.5 (E0.5). Embryos were collected at E9.5. Trunk neural plates were dissected adjacent to somites 18-21. Neural plate borders were then isolated producing two explants per embryo, which were then plated individually onto 1μg/ml fibronectin pre-coated glass bottomed 24-well dishes. Explants were cultured in neural crest conditioned media as described in Gonzalez Malagon et al. 2019 at 37°C at 5%CO_2_^30^.

### Molecular Biology and transfection

HEK293T cells, O9-1 cells and primary mouse trunk neural crest explants were transiently transfected using Lipofectamine 2000 (Invitrogen, 11668019) according to manufacturer’s instructions. For primary trunk neural crest explants, cells were transfected 24h post-dissection. Following plasmids were used: PB_EF1α_eGFP or PB_EF1α_ALK^WT/F1174L/R1275Q^-eGFP for generating transgenic O9-1 cell lines, pCAG_eGFP or pCAG_ALK^WT/F1174L/R1275Q-^eGFP for live-imaging, pCAG_GSK3α/β, pCAG_GSK3α/β_Flag/Strep, pCAG_ALK^WT/F1174L/R1275Q/I1250T^-Flag/Strep, pcDNA3.1_ALK^WT/F1174L/R1275Q^ for immunoprecipitation assays.

For detailed plasmid generation and transfection protocols, see Supplementary Methods.

### Pharmacological inhibition

Crizotinib (LC labs, C-7900) was resuspended in dimethyl sulfoxide (DMSO) (Sigma-Aldrich, M81802) at a stock concentration of 10 mM and stored at −20°C. As control, DMSO was diluted to match the maximum amount of added stock solution. Crizotinib was diluted with complete HEK293T media for a final concentration of 0.5 μM and added to cells.

### Immunoprecipitations and Western blot analysis

For immunoprecipitations assays, cells were lysed using glutathione S-transferase (GST) buffer (50 mM Tris-HCl pH 7.4, 200 mM NaCl, 1% NP-40, 2 mM MgCl2, 10% glycerol) supplemented with cOmplete™ EDTA-free Protease Inhibitor Cocktail (Merck, 11836170001) and PhosSTOP™ (Merck, 4906845001) and performed using Anti-FLAG® M2 Magnetic Beads (Sigma, M8823) according to manufacturer’s instructions. 10μl of M2 Magnetic beads were incubated with 200μg protein.

Western blot analysis was done using XCell SureLock™ Mini-Cell and XCell II™ Blot Module Electrophoresis System (ThermoFisher, EI0002) according to manufacturer’s instructions. For total lysate, 10μg protein was added and 5 μl of Precision Plus Protein Dual Color Standards (BioRad, 1610374) as protein ladder.

All primary antibodies were diluted 1:1000; ALK (CST, 3633T), p(Y1507)ALK (CST, 14678), Flag (Sigma, F1804-200UG), GFP (CST, 2956), GSK3α (CST, 9338), GSK3β (CST, 9315), GSK3α/β (CST, 5676), HSP90 (Santa Cruz, SC-13119).

### Immunofluorescence for O9-1 cells and trunk neural crest cells

O9-1 cells were fixed with 4%PFA at room temperature and permeabilised with 0.4% Triton X100 in PBS for 10 min. Primary and secondary antibodies were incubated for 1 hour in in 3% BSA in PBS blocking buffer. Neural crest explants were fixed and stained as described in ^30^. Cells were mounted using Fluoroshield Mounting Medium with DAPI (Abcam, ab104139) and imaged at 20x magnification using a ZEISS LSM 980. Phalloidin IgG, Alexa Fluor 568 (Invitrogen, A12380) was incubated with secondary antibodies. Primary and secondary antibodies were used at 1:500 dilution: GFP (Abcam, ab13970), Ki67 (Invitrogen, 14-5698-82), Goat anti-chicken IgG, Alexa Flour 488 (Invitrogen, A-11039), Donkey anti-rat IgG, Alexa Flour 647 (Abcam, ab150155). Images were taken at the same exposure time and laser power across the same conditions and images were analysed using ImageJ. For cell circularity and area, ten cells per image were analysed.

### Live-cell imaging and quantification

Live cell imaging was done 24hr’s post-transfection with PhaseFocus at 4x magnification or with ZEISS LSM 980 at 10x magnification at 37°C with 5% CO_2_. Phase-contrast (on PhaseFocus) or transmitted light module (Zeiss) were used to image all migrating cells; a 488nm filter was used to detect GFP positive cells. Imaging was done over 24 hours, 20 min per frame. Per explants, a minimum of two regions were imaged.

GFP positive cells were manually tracked using the ImageJ plug in Manual Tracking. Cells were only tracked if they were present in three or more images and track was terminated when cells were dividing. Manual tracking generated XY coordinated with distance and migration velocity over time. XY coordinates were used to generate a migration rose plot and calculate direction autocorrelation using Matlab. This was done as describe in Gorelik & Gautreau, 2014^28^.

### Protein-Protein Structure Modelling and Molecular Dynamics Simulations

Crystal structures of ALK (UniProt: Q9UM73), GSK3α (UniProt: P49840), and GSK3β (UniProt: P49841) were retrieved from the UniProt database. AlphaFold was utilized to resolve missing or unresolved regions. Models were validated using Ramachandran plot analysis to ensure stereochemical accuracy, identifying regions requiring refinement. This process yielded reliable structural models for the eight pairwise systems: ALK (ALK, ALK^F1174L^, ALK^R1275Q^, ALK^I1250T^) versus GSK3 (GSK3α, GSK3β).

Protein-protein docking was performed using the High Ambiguity Driven Biomolecular Docking (HADDOCK) platform with default parameters. The complex with the lowest binding energy was selected for each of the eight systems for subsequent molecular dynamics simulations.

Molecular dynamics (MD) simulations were conducted for each of the eight ALK-GSK3 complexes using GROMACS (version 2022). MD simulations were run for 50 ns with a 2fs time step, employing the LINCS algorithm to constrain hydrogen bonds. Trajectories were saved every 10 ps for analysis. Using GROMACS tools, Root mean square deviation (RMSD), root mean square fluctuation (RMSF), and radius of gyration (Rg) were calculated. Simulation trajectories were visualized using Visual Molecular Dynamics (VMD), Moludock Molecular Viewer (https://lutimbastuart.github.io/Molecular_Carousel/) and UCSF Chimera.

Specific interaction sites and key residues mediating ALK-GSK3 interactions were identified using the Protein-Ligand Interaction Profiler (PLIP) and visualized with PyMOL. Intermolecular contacts, including hydrogen bonds, hydrophobic interactions, and salt bridges, were quantified across the simulation trajectories to characterize the binding interfaces of the eight complexes. The raw data files used and generated in the molecular dynamic simulation analysis have been deposited in a GitHub repository link accessible at: https://github.com/lutimbastuart/ALK_Variants_In_Neuroblastoma.

See Supplementary Methods for more detailed ALK-GSK3 modelling.

## Acknowledgements

We thank members of the Liu lab, the Centre for Craniofacial and Regenerative Biology and King’s College London Biological Services Unit for their support. We thank labs of Matthias Krause and Louis Chesler for reagents and support. Core funding from London South Bank University for establishment of computational biology laboratory (MoLuDock Lab, MAM, SL). This study was funded by grants from UKRI Medical Research Council MC_PC_21044 (KJL, AMW, JD) and the European Union’s Horizon 2020 research and innovation programme, Marie Skłodowska-Curie grant agreement No 860635 (AMW, KJL), and a collaborative MRC Doctoral Training Programme iCASE studentship with STEMCELL Technologies (DC, KJL). Results above reflect only the author’s view and that the Research European Agency is not responsible for any use that may be made of the information it contains.

