## Supplemental figures and methods for "Neuroblastoma-associated ALK variants have distinct cellular and biochemical activities"

**This PDF file includes:**

Supporting Text
Figures S1
Legends for Movies S1 to S3
SI Methods
SI References

**Other supporting materials for this manuscript include the following:**

Movies S1 to S3

**Supporting Text**

<u>Advanced structural modeling of ALK with GSK3 isoforms</u>

To investigate how specific ALK mutations influence the structural dynamics of its complexes with GSK3, we performed all-atom molecular dynamics (MD) simulations of ALK bound to both GSK3 isoforms (α and β), examining the wild-type (WT) protein alongside the F1174L, R1275Q, and kinase-dead I1250T variants.

The wildtype ALK kinase domain (residues 1105-1377) was observed to interact with the kinase domain of GSK3α (residues 82 to 457) in a protein-protein complex as inferred from the modelling approach (Figure 4A). Both proteins exhibit a characteristic bilobal kinase architecture with interactions occuring at a defined cleft between the ALK and GSK3α, and a protein-protein interaction (PPI) distance of ≤3Å, highlighting close contacts (figure 4A). For ALK-GSK3α complex key amino acids involved in the interaction are for ALK: His1124, Glu1154, Glu1158, Glu1129, Tyr1282, Cys1288 and for GSK3α Arg56, Arg93, Arg95, Asn335, Arg336, Phe344 (Figure 4A). Interaction is primarily facilitated through polar or charged amino acids (Glu, Arg, His) with contributions through hydrogen bonds, hydrophobic interaction, and possibly π-π stacking via Phe334 and Tyr1282. On the other hand, the interaction between ALKWT and GSK3β is facilitated through flexibility of the interface and hydrophobic contacts (Phe1174, Val1180) with interacting amino acids for ALK: Ile1095, Lys1109, Lys1101, Asn1140, Ile1171, Phe1174, Gln1177, Val1180, Arg1181, Gln1238, Val1265, Val1268, Ala1377, Gly1596, and for GSK3β Met1, Ser9, Arg223, Thr232, Val263, Gln287 (Figure 4B). Although ALK interacts with GSK3α and GSK3β in slightly different ways, their binding pockets are largely the same. The two isoforms have very similar sequences, but small shifts in their kinase lobes especially the N-terminus, which may influence their specificity and interaction strength with partners like ALK.

<u>The F1174L Mutation Compromises Complex Stability.</u>

At the interface, flexibility from this region allows close contact with ALK, facilitating both hydrophobic stacking promoting more robust and adaptable protein–protein interaction (Figure 4B). GSK3β's greater conformational flexibility at its N-lobe allows altered cleft closure with ALK, engaging more hydrophobic contacts (Phe1174, Val1180). GSK3α complex, in contrast, tend toward a more open cleft with fewer hydrophobic and more polar contacts, attributed to subtle residue differences and local loop rigidity (figure 4A/B). Although ALK interacts with GSK3α and GSK3β in slightly different ways, their binding pockets are largely the same. The two isoforms have very similar sequences, but small shifts in their kinase lobes especially the N-terminus which may influence their specificity and interaction strength with partners like ALK[1,2].

To assess the global stability and conformational drift of the ALK-GSK3 complexes, we calculated the Root Mean Square Deviation (RMSD) of the protein backbone over a 50 ns simulation trajectory. Notably, the F1174L mutant complex exhibited the highest average RMSD in both isoforms (GSK3α: 6.26 ± 2.76 Å; GSK3β: 7.39 ± 2.46 Å) (Figure 4C/D). This significant increase in structural deviation, marked by a larger standard deviation, indicates a more dynamic and less stable complex conformation compared to WT (GSK3α: 3.82 ± 1.07 Å; GSK3β: 5.06 ± 1.30 Å). Primarily, these findings are consistent with recent literature demonstrating that increased protein flexibility reflected by higher RMSD can foster alternative, functionally relevant conformational states, ultimately enhancing allosteric regulation and enzymatic output rather than merely weakening binding affinity[3]. In stark contrast, the R1275Q mutant complex with GSK3α was remarkably stable (3.70 ± 1.02 Å), mirroring the WT. Interestingly, the R1275Q mutation induced pronounced destabilization in the GSK3β complex (8.19 ± 2.47 Å), suggesting isoform-specific induced structural impacts. The kinase-dead I1250T mutant displayed intermediate destabilization.

We next probed local flexibility and structural compactness using Root Mean Square Fluctuation (RMSF) and Radius of Gyration (Rg) analyses. The RMSF analysis revealed that all mutations increased local residue fluctuations relative to WT (Figure 4E/F), with the I1250T mutant showing the highest per-residue flexibility. Critically, the F1174L mutation induced substantial fluctuations across the complex, particularly in the GSK3β isoform (9.99 ± 2.62 Å vs. WT 5.88 ± 1.50 Å). This pronounced flexibility translated into a measurable expansion of the complex. The Rg analysis confirmed that the F1174L mutant yielded the most expanded conformational ensemble for the GSK3β complex (29.86 ± 0.59 Å vs. WT 28.58 ± 0.37 Å) (Figure 4H). A similar, though less dramatic, expansion trend was observed for F1174L in the GSK3α complex (29.04 ± 0.41 Å vs. WT 28.78 ± 0.36 Å) (Figure 4G). This mutation-induced loosening of the complex architecture stands in clear contrast to the R1275Q mutant, which had a minimal effect on the compactness of the GSK3α complex.

ALK-GSK3 Binding Interfaces across Oncogenic Mutants

With reference to the supplementary figure S1, in the $ALK^{F1174L}$-GSK3α complex (Supplementary Figure S1A), the contact surface is dominated by extensive hydrophobic and polar interactions involving for $ALK^{F1174L}$: Ala1126, Leu1190, Gln1287, Trp1295, Pro1298 His1368, Gln1367, and GSK3α: Leu144, Tyr197, Gln412, Pro418, , Ser447, Ala449, Leu454. The interface benefits from arginine–rich patches (Arg417, Arg142, Arg176), known to stabilize transient electrostatic contacts. This interaction pattern suggests both robustness and local adaptability, consistent with the F1174L-driven conformational dynamics and complemented by prior findings showing that increased flexibility at kinase interfaces can facilitate allosteric transitions and functionally relevant complexation. The repeated appearance of residues such as Leu454 and Gln412—engaged from multiple domains may indicate intramolecular bridging.

The $ALK^{F1174L}$-GSK3β complex (Supplementary Figure S1D) introduces a distinct set of interacting residues for $ALK^{F1174L}$: Lys1101, Leu1174, Leu1224, Glu1242, Pro1357, Val1358. Phe1376 and GSK3β: Arg6, Thr8, Arg92, Gly230, Thr232, Asn287, Tyr288, Phe291, Lys292. Here, the presence of several small polar and basic residues at the interface (Thr8, Arg6, Asn287, Lys292) highlights the likely involvement of flexible loop elements particularly around the N-lobe and glycine-rich regions.

In the ALK$^{R1275Q}$-GSK3α complex (Supplementary Figure S1B), the interface pivots for ALK$^{R1275Q}$ on: Leu1190, Glm1275, Tyr1282, Glu1340, Thr1343, and for GSK3α on: Lys11, Glu59, Arg61, Lys114, Asp172, Thr175, Arg326, Arg336, Phe342, Arg365, Pro375, Thr374. The inclusion of the mutant residue Glu1275 and the maintenance of several stabilizing salt-bridges (Lys114, Glu59, Arg661, Glu1340, Thr1343) suggest that, while the mutant partly retains the capacity for wild-type-like interaction, the subtle alteration of charge and hydrogen-bonding potential may underlie the intermediate dynamics seen in simulation and partial activity retention observed in functional assays[5]. With reference to the ALK$^{R1275Q}$–GSK3β complex (Supplementary Figure S1E), the following amino acids mediate the interaction for ALK$^{R1275Q}$: Lys1097, Ala1099, Gly1100, Arg1231, His1244, Gln1275, and GSK3β: Arg4, Arg6, Thr7, Lys15, Glu137, Ser215, Phe293. This pattern combines wild-type residue contacts (Gly1100, Thr7, Ala1099) with uniquely mutant interactions (Gln1275), along with a mixture of aromatic and charged residues facilitating a flexible yet functionally capable interface. The preservation of several positively charged residues and exposure of mutant side chains is consistent with the slightly enhanced but less robust interaction models for ALK$^{R1275Q}$.

In the ALK$^{I1250T}$-GSK3α complex (Supplementary Figure S1C), key contacts include for ALK$^{I1250T}$: Val10, Arg1120, Phe1127, Glu1129, Tyr1131, Leu1190 , Thr1250, Arg1284, Val1293, Phe1301, and GSK3α: Arg56, Arg61, Glu62, Leu325, Lys327, Pro339, Thr371, Thr372. Notably, extensive arginine and glutamate clustering (Arg61, Arg56, Arg1120, Glu62, Glu1129) suggests persistent charge complementarity, but the presence of noncanonical residues at the interface, and increased involvement of flexible termini (Val10, Thr371, Thr372), likely disrupt the ordered hydrogen bond and hydrophobic networks seen in wild-type state. This faulty engagement pattern recapitulates observations that broken or misaligned contacts in kinase-dead or inactivating mutants result in instability and loss of proper conformational coupling[4].

For ALK$^{I1250T}$ with GSK3β (Supplementary Figure S1F) amino acids involved in the interaction are for ALK$^{I1250T}$: Asp1110, Pro1112, Phe1174, His1176,

Leu1227, Asp1238.Thr1275 and for GSK3β: Arg4, Thr8, Phe93, Leu122, Asp125, Pro212, Asn213, Gly230, Asn285, Asn287. This constellation includes flexible loop and C-terminal elements (Asn287, Leu1227, Thr1275), charged and aromatic side chains, and critical mutants (Phe1174, His1176), closely mirroring the expanded dynamics and instability inferred from in silico metrics and previous kinase functional analyses. The apparent dispersal of interactions across the GSK3β sequence may reflect compensatory stabilization due to the loss of canonical interface geometry.

Through molecular modelling approaches, we revealed specific residue-level changes at the ALK-GSK3 interface directly govern global conformational dynamics. Our analyses demonstrate that the F1174L mutation facilitates a plasticity-enhancing interface, the I1250T kinase-dead mutation disrupts functional complex geometry, and the R1275Q mutation exerts critically isoform-dependent effects, thereby elucidating a key mechanistic basis for differential signaling in neuroblastoma disease development.

172 **Figures**

173

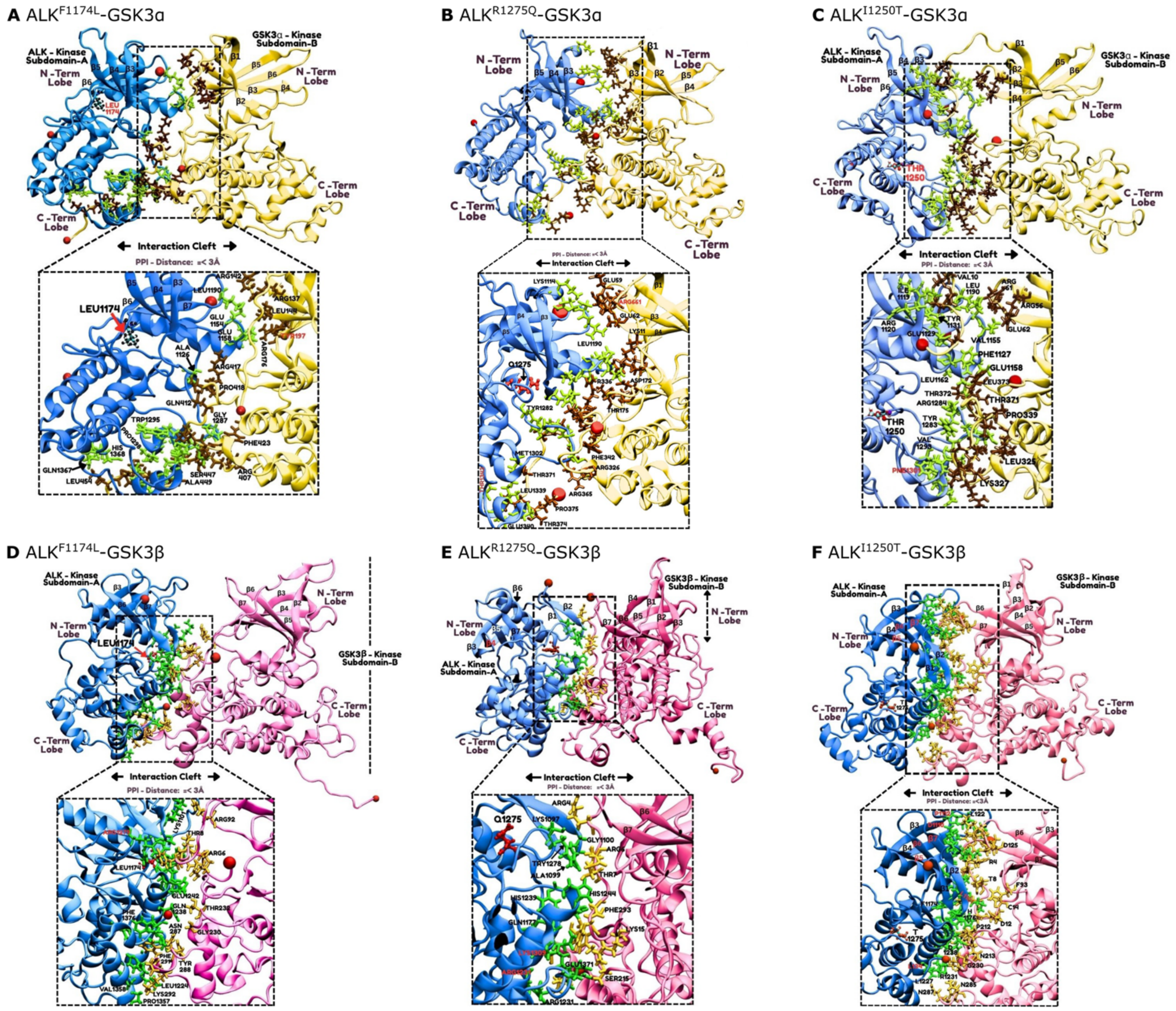
A ALK$^{F1174L}$-GSK3α
B ALK$^{R1275Q}$-GSK3α
C ALK$^{I1250T}$-GSK3α
D ALK$^{F1174L}$-GSK3β
E ALK$^{R1275Q}$-GSK3β
F ALK$^{I1250T}$-GSK3β
ALK - Kinase Subdomain-A
GSK3α - Kinase Subdomain-B
GSK3β - Kinase Subdomain-B
N -Term Lobe
C -Term Lobe
Interaction Cleft
PPI - Distance: =< 3Å
LEU1174
Q1275
THR 1250

174

**Figure S1**: **Structural representation of protein–protein interfaces between ALK mutants and GSK3 isoforms obtained from 50 ns molecular dynamics simulations. (Associated with Figure 4.)**

GSK3α (yellow) interaction with ALK mutants (blue): (A) ALK$^{F1174L}$, (B) ALK$^{R1275Q}$, (C) ALK$^{I1250T}$. GSK3β (pink) interaction with ALK mutants (blue): (D) ALK$^{F1174L}$, (E) ALK$^{R1275Q}$, (F) ALK$^{I1250T}$. Enlarged interaction cleft (distance <3 Å) reveals side-chain contacts that stabilize the complex: ALK residues are shown as green sticks, and GSK3 residues in brown (α) and yellow (β).

**Movie S1: ALK overexpression in primary trunk neural crest cells leads to increased migration and loss of directionality. (Associated with Figure 2.)**

Primary trunk neural crest explants from E9.5 mouse embryos were transfected 24 hours post-dissection. Post-transfection, 24 hour time lapse imaging of the migratory trunk neural crest cells was performed (20min/1 frame) using PhaseFocus. Cells migrate from the neural plate border (left) toward right. Movies are linked to still images in Figure 2E. eGFP cells were manually tracked using ImageJ (example track blue). Scale bar: 100μm

**Movie S2: Molecular dynamic simulation of ALK-GSK3α complex. (Associated with Figure 4.)**

50 ns molecular dynamic simulation of GSK3α (Red) in complex with either $ALK^{WT}$, $ALK^{F1174L}$, $ALK^{R1275Q}$, or $ALK^{I1250T}$ (Cyan). Movie trajectories were analyzed to infer structural dynamics visualized through structural metrics such as root mean square deviation (RMSD), root mean square fluctuation (RMSF) and radius of gyration (Rg) as reported in Figure 4C/E/G. Visualizations were rendered at 0.2ns/frame. Amino acids involved in the interaction are depicted as Ball-and-Stick. Mutations are shown as sticks.

**Movie S3: Molecular Dynamic simulation of ALK-GSK3β complex. (Associated with Figure 4.)**

50 ns molecular dynamic simulation of GSK3β (red) in complex with either $ALK^{WT}$, $ALK^{F1174L}$, $ALK^{R1275Q}$, or $ALK^{I1250T}$ (Cyan). Movies trajectories were analyzed to infer structural dynamics, visualized through structural metrics such as root mean square deviation (RMSD), root mean square fluctuation (RMSF) and radius of gyration (Rg) in Figure 4D/F/H. Visualizations were rendered at 0.2ns/frame. Amino acids involved in the interaction are depicted as Ball-and-Stick. Mutations are shown as sticks.

**SI Methods**

Molecular Biology and transfection

For generating plasmids the following materials were used: PB-Ef1α-Tet3G-T2ANeo (kindly donated by Andrea Serio lab), pcDNA3.1_ALK$^{WT}$, pcDNA3.1_ALK$^{F1174L}$, pcDNA3.1_ALK$^{R1275Q}$, pCAG_Dest_Flag/Strep (kindly donated by Matthias Kause lab), pCAG_Dest_eGFP (kindly donated by Matthias Krause lab), pEntry_ALK$^{WT}$ (Addgene 23917), pDONR221_GSK3α (DNASU, HsCD00039881), pDONR221_GSK3β (DNASU, HsCD00041464), pcs22424_GSK3α (original plasmid from Addgene, 22424), pcs22424_GSK3β (original plasmid from Addgene, 22424).

pEntry_ALK$^{F1174L}$ and pEntry_ALK$^{R1275Q}$ were generating by restriction digest. In brief, pcDNA3.1_ALK$^{F1174L}$, pcDNA3.1_ALK$^{R1275Q}$ and pEntry_ALK$^{WT}$ were digested with BsmI (NEB, R0134S) and fragments containing the mutation site were ligated into the pEntry_ALK$^{WT}$ backbone using T4 ligase (Promega, M1801). pEntry_ALK$^{I1250T}$ was generated using site directed mutagenesis.

Gateway cloning LR clonase reaction (Thermo-Fisher, 11791020) was used to generate pCAG_ALK$^{WT/F1174L/R1275Q/I1250T}$_eGFP, pCAG_ALK$^{WT/F1174L/R1275Q/I1250T}$_Flag/Strep, pCAG_GSK3α/β_Flag/Strep by recombinating pCAG_Dest_eGFP or pCAG_Dest_Flag/Strep with pEntry_ALK$^{WT/F1174L/R1275Q/I1250T}$ or pDONR221_GSK3α/β.

pCAG_GSK3α and pCAG_GSK3β were generated by cutting full-length GSK3α and GSK3β from pcs22424_GSK3α) or pcs22424_GSK3β using EcoRI (Promega R601A) and NotI (Promega, R643A) and ligating it with pCAG_eGFP backbone using T4 ligase.

PB_EF1α_ALK$^{WT/F1174L/R1275Q}$-eGFP were generated using Gibson Assembly (NEB, E2611) by amplifying ALK$^{WT/F1174L/R1275Q}$-eGFP from pCAG_ALK$^{WT/F1174L/R1275Q}$-eGFP using PCR and assembling it with PB_Efα_T2A_Neo.

HEK293T cells, O9-1 cells and primary mouse trunk neural crest explants were transiently transfected using Lipofectamine 2000 (Invitrogen, 11668019) according to manufacturer's instructions. For HEK293T cells, 1x106 HEK293T cells were plated and 24hr's later, cells were transfected with 5µl of Lipofectamine 2000 and a total of 2µg of plasmid. For O9-1 cells, 1.3x104 cells were plated and 24hr's later, cells were transfected with 2µl Lipofectamine 2000 and a total of 2µg of plasmid. For primary trunk neural crest explants, cells were transfected with 2µl of Lipofectamine 2000 and a total of 0.5µg of plasmid 24h post-dissection.

Protein Structure Acquisition and Modelling

Crystal structures of ALK (UniProt: Q9UM73), GSK3α (UniProt: P49840), and GSK3β (UniProt: P49841) were retrieved from the UniProt database. AlphaFold was utilized to resolve missing or unresolved regions. Models were validated using Ramachandran plot analysis to ensure stereochemical accuracy, identifying regions requiring refinement. This process yielded reliable structural models for the eight systems: $ALK^{WT}$-GSK3α, $ALK^{WT}$-GSK3β, $ALK^{F1174L}$-GSK3α, $ALK^{F1174L}$-GSK3β, $ALK^{R1275Q}$-GSK3α, $ALK^{R1275Q}$-GSK3β, $ALK^{I1250T}$-GSK3α, $ALK^{I1250T}$-GSK3β complexes.

Protein-Protein Docking

Protein-protein docking was performed using the High Ambiguity Driven biomolecular Docking (HADDOCK) platform with default parameters. The docking process involved three stages: (1) rigid-body docking to generate a large ensemble of potential complex structures, (2) semi-flexible simulated annealing molecular dynamics to refine these structures, and (3) free energy calculations to rank the refined structures based on binding energies. The complex with the lowest binding energy was selected for each of the eight systems for subsequent molecular dynamics simulations.

Molecular Dynamics Simulations

Molecular dynamics (MD) simulations were conducted for each of the eight ALK-GSK3 complexes using GROMACS (version 2022). The systems were parameterized with the CHARMM36 force field and solvated in a cubic box of TIP3P water molecules, with $Na^{+}$ and $Cl^{-}$ ions added to neutralize the system and achieve a physiological salt concentration of 0.15 M. Each system underwent energy minimization using the steepest descent algorithm, followed by equilibration in the NVT (constant number of particles, volume, and temperature) and NPT (constant number of particles, pressure, and temperature) ensembles for 100 ps each at 310 K and 1 bar. Production MD simulations were run for 50 ns with a 2 fs time step, employing the LINCS algorithm to constrain hydrogen bonds. Trajectories were saved every 10 ps for analysis.

Conformational Analysis

Root mean square deviation (RMSD), root mean square fluctuation (RMSF), and radius of gyration (Rg) were calculated using GROMACS tools. RMSD was computed for Cα atoms relative to the initial structure after least-squares fitting, evaluating deviations in the bound and apo states over the simulation time. RMSF was calculated for Cα atoms to quantify per-residue flexibility, with the average structure derived from simulation snapshots. Rg was determined to assess the compactness of each complex.

Interaction Analysis

Specific interaction sites and key residues mediating ALK-GSK3 interactions were identified using the Protein-Ligand Interaction Profiler (PLIP) and visualized with PyMOL. Intermolecular contacts, including hydrogen bonds, hydrophobic interactions, and salt bridges, were quantified across the simulation trajectories to characterize the binding interfaces of the eight complexes.

Visualization and Rendering

Simulation trajectories were visualized using Visual Molecular Dynamics (VMD), Moludock Molecular Viewer and UCSF Chimera. VMD was employed to load DCD trajectory files, align frames to the reference structure using the RMSD tool, and visualize binding pockets with the Volume Viewer tool. Animations of the trajectories were generated to observe dynamic interactions. Chimera was used to create 3D representations of protein surfaces and binding modes, with the Surface tool for surface rendering and the Surface Color tool to highlight key regions, including binding pockets. These tools facilitated detailed analysis of protein-protein interactions and conformational dynamics.
